# Modality-specific dysfunctional neural processing of social-abstract and non-social-concrete information in schizophrenia

**DOI:** 10.1101/2020.02.18.953927

**Authors:** Yifei He, Miriam Steines, Gebhard Sammer, Arne Nagels, Tilo Kircher, Benjamin Straube

## Abstract

Schizophrenia is characterized by marked communication dysfunctions encompassing potential impairments in the processing of social-abstract and non-social-concrete information, especially in everyday situations where multiple modalities are present in the form of speech and gesture. To date, the neurobiological basis of these deficits remains elusive. In a functional magnetic resonance imaging (fMRI) study, 17 patients with schizophrenia or schizoaffective disorder, and 18 matched controls watched videos of an actor speaking, gesturing (unimodal), and both speaking and gesturing (bimodal) about social or non-social events in a naturalistic way. Participants were asked to judge whether each video contains person-related (social) or object-related (non-social) information. When processing social-abstract content, patients showed reduced activation in the medial prefrontal cortex (mPFC) only in the gesture but not in the speech condition. For non-social-concrete content, remarkably, reduced neural activation for patients in the left postcentral gyrus and the right insula was observed only in the speech condition. Moreover, in the bimodal conditions, patients displayed improved task performance and comparable activation to controls in both social and non-social content. To conclude, patients with schizophrenia displayed modality-specific aberrant neural processing of social and non-social information, which is not present for the bimodal conditions. This finding provides novel insights into dysfunctional multimodal communication in schizophrenia, and may have potential therapeutic implications.

## 1. Introduction

In everyday social communication, patients with schizophrenia encounter a diverse spectrum of inputs from multiple modalities (Holler & Levinson, 2019). These include non-linguistic stimuli from others’ facial expressions (Gur et al., 2002; Vuilleumier et al., 2001), body movements, e.g., postures and gestures (He et al., 2020; Nagels et al., 2015b; Straube et al., 2013a), and linguistic stimuli in the form of auditory speech and written texts (Brown & Kuperberg, 2015). Importantly, the multimodal inputs comprise both social-abstract and non-social-concrete information, and patients’ social functioning heavily depends on the processing of these stimuli, which serves as the basis to further mentalize social intentions and to perform appropriate social interaction (Amodio & Frith, 2006). To date, however, it remains elusive if the processing of these social/non-social information in schizophrenia is impaired in schizophrenia in a multimodal context.

Social cognition is known to be impaired in schizophrenia (Green et al., 2015). Regarding the perception of social stimuli, in particular, prior research on patients’ social perception dysfunction has focused on emotional perception of faces and voices (Anticevic et al., 2012; Witteman et al., 2012), whereas limited studies have investigated the perception of social-abstract information delivered via linguistic and gesture stimuli. In healthy participants, a seminal fMRI (functional magnetic resonance imaging) study has identified distinct brain regions for processing socially and non-socially relevant linguistic information (Mitchell et al., 2002): When participants were asked to judge whether visual word pairs are person– or object-related, person-related social stimuli activated the medial prefrontal cortex (mPFC), a crucial region forming the mentalizing network (Amodio & Frith, 2006; Frith & Frith, 2006; Saxe & Powell, 2006; Van Overwalle, 2009). On the other hand, when processing linguistic stimuli about non-social-concrete content (e.g., words and sentences about objects), healthy participants activate regions including a left-lateralized network including the left the bilateral insula and the left parietal lobe (Binder et al., 2005; Chao et al., 1999).

Notably, in everyday life, both types of information are also commonly conveyed via non-linguistic channels such as manual gestures. For instance, even without speech, individuals can use a “be silent” emblematic gesture to deliver social information, and a “hammering” pantomime to describe a concrete object or action. Moreover, it is common to use both gesture and speech together: To ask someone else to stop, one can use a “stop” gesture (e.g., a front-facing, raised hand) together with its verbal counterpart. Emerging basic research has investigated if the processing of social-abstract and non-social-concrete information is modality-independent: for healthy populations, social-abstract information delivered by both auditory speech and manual gesture commonly activates the mPFC and the (IFG) and middle temporal gyri (Straube et al., 2010; Straube et al., 2013b)—regions that are typically activated for social cognition and abstract semantics (Binder et al., 2009; Van Overwalle, 2009). For processing non-social action and object information, literature also suggests that humans recruit modality-independent regions, such as the lateral occipitotemporal cortex (LOTC) and the pre/postcentral gyri (Barsalou, 2008; Pulvermüller et al., 2005; Wurm & Caramazza, 2019).

The multimodal characteristics of social/non-social information processing, as well as its supramodal neural basis, may have profound implications in schizophrenia research. Schizophrenia is characterized by well-known deficits in the comprehension and production of speech (Bleuler, 1950; Kircher et al., 2002; Kuperberg et al., 2008), as well as impairments in the perception and production of gesture (Matthews et al., 2013; Millman et al., 2014; Mittal et al., 2006; Nagels et al., 2018; Straube et al., 2014; Straube et al., 2013a; Thakkar et al., 2014; Walther et al., 2015; Walther et al., 2013). Both language and gesture dysfunctions in schizophrenia are indicative of negative and positive symptoms of schizophrenia (Park et al., 2008; Walther et al., 2019), and are functionally highly relevant (Friedman et al., 2012; Walther et al., 2016). Yet, regarding the processing of social/non-social information, it remains unknown if both processes are impaired in schizophrenia irrespective of modality, or if the impairments are modality-specific. For the processing of social-abstract information, both linguistic (visual and auditory) and gesture (social emblems) have not been investigated in schizophrenia so far. However, studies employing non-semantic hand actions have shown that patients might be impaired for gesture perception in general (Thakkar et al., 2014; Walther et al., 2015). For the processing of concrete and non-social information, a previous study shows that patients are impaired in visually presented sentences (Kuperberg et al., 2008). Moreover, given potential dysfunctional processing of social or non-social information in schizophrenia, it remains unclear if these deficits could be potentially compensated by multimodal inputs containing both speech and gesture: Emerging literature suggests mutual facilitation between speech and gesture, at least for healthy populations (Cuevas et al., 2019; Drijvers et al., 2018; He et al., 2018b; Krahmer & Swerts, 2007; Wang & Chu, 2013).

To address these remaining research questions, we conducted the current study, presenting to patients with schizophrenia and matched controls with videos of an actor communicating in a spontaneous and naturalistic manner. In these videos, the actor performs either social-abstract (person-related) or non-social (object-related) content in different modalities, where social and non-social information is perceivable in gesture- and speech-only modalities. Similar to approaches from previous research (Mitchell et al., 2002; Straube et al., 2013b), we directly compared social vs. non-social videos to identify neural perception of social and non-social information in both auditory-speech and visual-gesture modalities. Additionally, we showed to participants videos with bimodal inputs (actor both speaking and gesturing). Based on previous research, we hypothesized activation of the mPFC and a left frontal-temporal network (e.g., inferior frontal gyrus, middle temporal gyrus) for the processing of social-abstract information (Mitchell et al., 2002). For non-social-concrete information processing, we hypothesized left-lateralized regions including the lateral occipitotemporal cortex (LOTC), the superior temporal gyrus/sulcus (STG/STS), as well as pre/postcentral gyri forming the putative mirror neuron system (Järveläinen et al., 2004; Johnson-Frey, 2004; Johnson-Frey et al., 2003; Lingnau & Downing, 2015). We focused directly on group differences between a group of patients suffering from schizophrenia or schizoaffective disorder, and their age- and education-matched controls: for social content, as patients are well-known for their social cognition impairments, we expected patients to show reduced activation in the mPFC, irrespective of encoding modality (Frith, 2004); For non-social content, despite mixed findings from previous neuroimaging research on hand action observation on schizophrenia (Horan et al., 2014; Thakkar et al., 2014), following previous report on dysfunctional processing of non-social linguistic stimuli in schizophrenia (Kuperberg et al., 2008), we hypothesized neural modulation of the object-related regions for patients with schizophrenia for both gesture and speech modalities. Additionally, based on prior basic research on mutual facilitation between gesture and speech (Cuevas et al., 2019; see Holler & Levinson, 2019 for review), we hypothesized that the bimodal input could compensate for potential unimodal processing deficits, leading to improved performance in the patient group.

## 2. Methods

### 2.1 Participants

We summarized participants’ demographic and clinical characteristics in **Table 1**. Healthy controls were recruited matching age and education to patients (Sassenhagen & Alday, 2016). Seventeen patients were recruited at the Department of Psychiatry and Psychotherapy at the Philipps University of Marburg, and were diagnosed according to ICD-10 with schizophrenia (F20.0, n=13, and F20.3, n=1) or schizoaffective disorder (F25.0, n=2, and F25.3, n=1). Participants in both groups are native speakers of German, and have no knowledge of Russian language. All except one of the patients received antipsychotic treatment; six were additionally treated with antidepressive medication. Positive and negative symptoms were assessed with the Scale for the Assessment of Positive Symptoms (SAPS) (Andreasen, 1984), and the Scale for the Assessment of Negative Symptoms (SANS) (Andreasen, 1981). Eighteen age- and education-matched healthy participants with no history of any mental disorders were recruited from the same area. Exclusion criteria for both groups were brain injury and neurological or other medical diseases affected by brain physiology. In both groups, we conducted neuropsychological tests to assess working memory function, digit span, trail making (TMT), verbal IQ (MWT-B) (Lehrl, 1999), and metaphoric language processing (concretism, evaluated with the Proverb Interpretation Task) (Barth & Küfferle, 2001). These measures are reported in **Table 1**. We report, additionally, scores from the subscales of SAPS and SANS (Andreasen, 1981, 1984), word fluency test, as well as gesture production and perception (BAG, Brief Assessment of Gesture (Nagels et al., 2015a)) in the supplement (**Table S1**). All participants had normal or corrected-to-normal vision and hearing. Except for one control and one patient, all other participants are right-handed (Oldfield, 1971). All participants gave written informed consent prior to participation in the experiment and were compensated monetarily. The study was approved by the ethics committee of the School of Medicine, Philipps University Marburg.

**Table 1.** Demographic, medication, symptom, and neuropsychological measures.

|  | Patients (n=17) | Controls (n=18) |
| --- | --- | --- |
| Age (years) | 33.12 (12.35) | 31.94 (10.21) |
| Gender male/female | 13/4 | 13/5 |
| Education (years) | 11.82 (1.77) | 12.72 (1.36) |
| TMT A (seconds) | 31.49 (10.73) | 26.17 (9.89) |
| TMT B (seconds) | 68.56 (37.8) | 52.93 (19.58) |
| Digit Span forward | 7.94 (1.75) | 8.05 (2.43) |
| Digit Span backward | 6.35 (1.93) | 6.61 (2.50) |
| Verbal IQ | 28.8 (5.25) | 28.5 (3.79) |
| *Concretism | 1.38 (0.45) | 1.14 (0.19) |
| SAPS (global) | 15 (6.89) |  |
| SANS (global) | 9 (6.02) |  |
| CPZ Equivalent | 562.52 (372.63) |  |

Values are presented as mean (SD). TMT: trail making test; CPZ: chlorpromazine. We reported only significant difference between controls and patients ( $p < 0.05$ , two-tailed t-test) with asterisks.

### 2.2 Materials and procedure

We employed a content judgement paradigm from previous studies from our research group to investigate modality-specific processing of social/non-social information (He et al., 2015; Straube et al., 2013b; Straube et al., 2018). Of note, the same fMRI dataset has been published with an unrelated research question (Wroblewski et al., 2020). We showed to participants five-second videos of an actor spontaneously communicating both social-abstract (S) and non-social-concrete (N) events in the following modalities: 1) incomprehensible Russian sentences with gestures. This is considered as a gesture-only (G) condition because social feature is only available to participants in the gesture form. 2) comprehensible German sentences (S) without any gestures. Additionally, we also showed to participants 3) German sentences with accompanying gestures as a bimodal input condition (B). A filler condition is also included with videos of incomprehensible Russian sentences with meaningless gestures. An example of both a social (S) and non-social (N) bimodal videos is illustrated in **Figure 1A**. For a complete list of all videos, please refer to Appendix in He et al. (2015).

**Figure 1.**
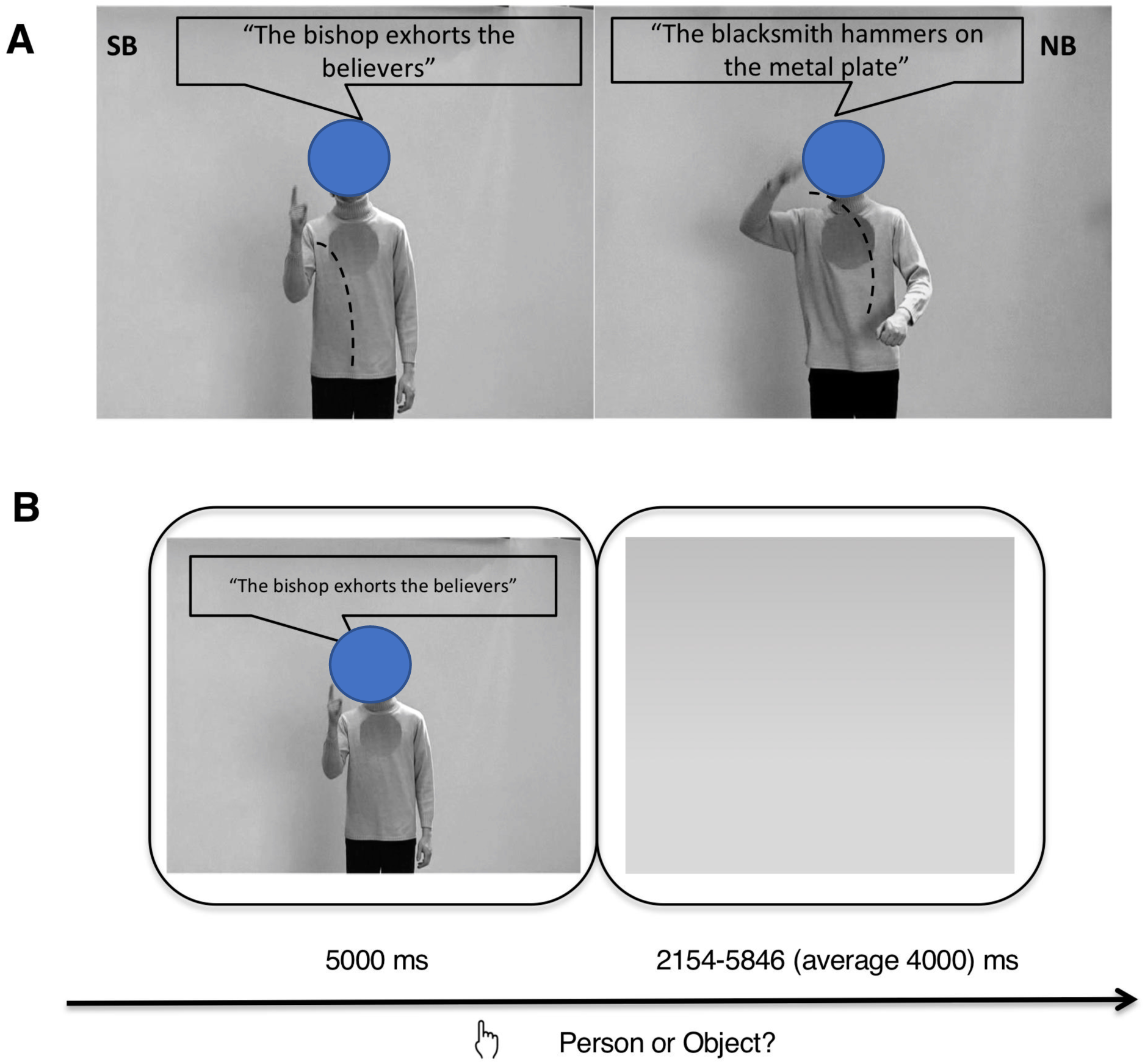
Panel A: Picture illustration for social-abstract (S) and non-social-concrete (N) videos in the bimodal condition (B). The same stimuli were also presented in two additional modalities: gestures with foreign Russian sentences (G) and German sentences without any gestures (S). For illustrative purposes, the spoken German sentences were translated into English, and all spoken sentences were written into speech bubbles. Panel B: Illustration of a sample trial. Participants performed a content judgment task for each video, indicating via button press whether a stimulus was either person– or object-related. Note that the face of the actor is only covered according to the privacy regulation from bioxariv.

### 2.3 Procedure

Altogether, 312 experimental video slips (26 videos per condition × 6 conditions × 2 sets) were included in the study. For each participant, an experimental session comprised 182 videos from one set of videos (156 critical videos and 26 filler videos), and consisted of two 14-minute runs. Each run contained 91 trials with a matched number of items from each condition. The stimuli were presented in an event-related design in pseudo-randomized order. Within each trial, each video-clip was followed by a gray background with a variable duration of 2154-5846ms (jitter average: 4000ms), as illustrated in **Figure 1B**. Participants performed a content judgement task for each video (Mitchell et al., 2002; Straube et al., 2013b), indicating via button press (with their left hand) whether a stimulus was either person– or object-related. Participants were instructed to respond to the task as soon as they had decided on an answer.

### 2.4 fMRI acquisition and preprocessing

All images were acquired using a 3T MRI scanner (Siemens MRT Trio series). The functional images were obtained using a T2*-weighted echo-planar image sequence (TR = 2s, TE = 30ms, flip angle = 90°, slice thickness = 4mm, interslice gap= 0.36mm, field of view= 230mm, matrix = 64 × 64, voxel size = 3.6 × 3.6 × 4.0mm, 30 axial slices orientated parallel to the AC-PC line, ascending order). Two runs of 425 volumes each were acquired during the experiment. Additionally, simultaneous EEG data from the participants were also collected for other analyses not relevant for the current study, and are therefore not further discussed here. MR images were preprocessed using the SPM12 software package (Statistical Parametric Mapping, Welcome Trust Center for Neuroimaging, London, UK) based on Matlab R2017a (version 9.2.0; MathWorks): after discarding the first five volumes to minimize T1-saturation effects, all images were spatially and temporally realigned, and normalized into the MNI space using the MNI template (resulting voxel size 2 × 2 × 2mm), smoothed (8mm isotropic Gaussian filter), and high-pass filtered (cut-off period 128s).

### 2.5 fMRI data analysis

We performed statistical whole-brain analysis in a two-level, mixed-effects procedure. On the first level, single-participant BOLD responses were modeled by a design matrix comprising the onset time points of each event (critical word of each sentence as used in the previous event-related fMRI and EEG studies, e.g., (He et al., 2015; He et al., 2018a; He et al., 2018c; Straube et al., 2013b; Straube et al., 2018)), with a duration of 5s for all experimental conditions. The micro-time onset was set to the average time bin (8 of 16) to align the onset vector to the slice in the middle of the brain. For all conditions, the duration of speech and gesture was used as parameters of no interests on a single trial level. Six movement regressors (three rotations and three translations) were entered in the single participant’s model to account for movement-induced effects on fMRI results. HRF was defined as the canonical HRF. Contrasts images against implicit baseline for all experimental conditions were used as summary measures and were included in the between-group analysis. We applied a flexible factorial analysis of variance using condition as main effect. We applied a Monte-Carlo simulation to determine the cluster extent threshold to correct for multiple comparisons, (Slotnick, 2017; Slotnick et al., 2003), which has been used in comparable studies in our laboratory on social perception of multimodal stimuli (Nagels et al., 2015b). For all statistical comparisons, the whole-brain activation was simulated assuming a voxel type-I error activation of p<.05, this revealed a cluster extent of 2268 contiguous resampled voxels as sufficient to correct for multiple comparisons at p<.0167 (Bonferroni-corrected for three independent tests for interaction within three independent modalities, with 0.05/3). We also reported in activation tables uncorrected cluster p-values and marked significance for cluster-level FWE correction. The reported voxel coordinates of activation peaks are located in MNI space. For the anatomical localization, functional data were referenced to the AAL toolbox (Tzourio-Mazoyer et al., 2002).

Firstly, we tested three-way interaction of group x modality x content. We then conducted, within each modality, interaction analyses to investigate group differences in the processing of social or non-social conditions, and masked the respective results based the contrast image from the first analysis, to reveal modality-specific group interaction. In addition, within each modality, for each group, we conducted pair-wise comparison between social and non-social conditions (S > N and N > S), for illustrating modality– and group-specific brain activations for either social or non-social information processing. In the end, following our hypotheses on bimodal enhancement for patients, within patients, we tested the interaction between modalities on social vs. non-social content processing, so as to reveal how bimodal stimuli might compensate potential neural processing deficits for patients with schizophrenia. Results of this analysis is reported in **Supplement S2**.

Based on the literature showing a potential relationship between symptom severity (especially negative symptoms) and social/non-social cognition (Mehta et al., 2014; Sergi et al., 2007), as well as gesture processing (Walther et al., 2019; Walther et al., 2013), for patients with schizophrenia, we conducted exploratory correlation analysis, probing for the potential relationship between clinical measures and brain activation in areas that are relevant to social/non-social information processing. To this end, spearman correlation analyses (uncorrected) were conducted between 1) parameter estimates from clusters showing significant group difference for either social or non-social conditions, 2) behavioral measures (reaction times and accuracy) for each experimental condition, and 3) scores from sum/general and subscales of SAPS and SANS.

## 3. Results

### 3.1 Behavioral results

Healthy controls and patients with schizophrenia were instructed to indicate via button press whether the actor in the video described a person-related content or an object-related content. Correct responses (percentage correct) and their reaction times were analyzed each with mixed ANOVA [within factors: CONTENT (social vs. non-social) and MODALITY (bimodal vs. gesture vs. speech); between factor: GROUP (control vs. patient)]. Descriptive statistics for each experimental condition for both groups were provided in **Table 2**.

**Table 2:** Accuracy and reaction times (RT) for the behavioral task

| **Accuracy (%)** |  |  |  |  |
| --- | --- | --- | --- | --- |
| **Conditions** | **Controls (n =18)** |  | **Patients (n = 17)** |  |
|  | **Mean** | **SD** | **Mean** | **SD** |
| Social Bimodal (SB) | 81.84 | 10.78 | 71.79 | 19.99 |
| Social Gesture (SG) | 70.29 | 16.36 | 63.34 | 18.45 |
| Social Speech (SS) | 79.91 | 17.17 | 71.71 | 20.57 |
| Non-social Bimodal (NB) | 93.95 | 6.19 | 91.18 | 12.96 |
| Non-social Gesture (NG) | 89.1 | 10.49 | 82.8 | 11.62 |
| Non-social Speech (NS) | 88.46 | 9.69 | 80.31 | 22.95 |

| **Reaction times (ms)** |  |  |  |  |
| --- | --- | --- | --- | --- |
| **Conditions** | **Controls (n =18)** |  | **Patients (n = 17)** |  |
|  | **Mean** | **SD** | **Mean** | **SD** |
| Social Bimodal (SB) | 3270.26 | 262.57 | 4196.31 | 831.15 |
| Social Gesture (SG) | 3154.67 | 281.27 | 4042.41 | 897.51 |
| Social Speech (SS) | 3455.9 | 279.19 | 4238.1 | 785.66 |
| Non-social Bimodal (NB) | 2982.74 | 262.79 | 4092.35 | 907.69 |
| Non-social Gesture (NG) | 3185.07 | 347.14 | 4122.33 | 924.68 |
| Non-social Speech (NS) | 3244.25 | 325.07 | 4177.06 | 921.39 |

SD, standard deviation; Reaction times were measured in reference to each video onset (full video length: 5s).

**Table 2:** Peak MNI coordinates of within- and between-group significant brain comparisons for social > non-social conditions. Cluster-level uncorrected p-valeue is provided for each cluster, asterisk indicates significance (p < 0.05) with cluster-level FWE correction.

| Contrasts | Anatomical Region (Peak) | Hem. | k | Peak MNI coordinates |  |  | t | p (unc.) |
| --- | --- | --- | --- | --- | --- | --- | --- | --- |
|  |  |  |  | x | y | z |  |  |
| Control |  |  |  |  |  |  |  |  |
| SS>NS | Inferior frontal gyrus pars orbitalis | L | 15202 | -48 | 22 | -6 | 6.74 | <.001* |
|  | Middle temporal gyrus | R | 2816 | 48 | -22 | -14 | 4.05 | .008 |
|  | Precuneus | L | 4169 | -4 | -48 | 42 | 3.18 | .002 |
| SG>NG | Superior frontal gyrus | L | 16869 | -22 | 52 | 32 | 4.68 | <.001* |
| SB>NB | Inferior frontal gyrus pars orbitalis | L | 9637 | -46 | 26 | -4 | 7.05 | <.001* |
|  | Supplementary motor area | L | 4916 | -8 | 16 | 66 | 5.3 | .001* |
|  | Middle temporal pole | R | 3023 | 50 | 10 | -28 | 4.64 | .006 |
| Patient |  |  |  |  |  |  |  |  |
| SS>NS | Inferior frontal gyrus pars orbitalis | L | 33204 | -50 | 20 | -4 | 5.77 | <.001* |
|  | Supplementary motor area | L | 7862 | -2 | 24 | 68 | 4.39 | <.001* |
| SG>NG | Superior frontal gyrus | L | 2836 | -16 | 32 | 62 | 3.86 | .008 |
| SB>NB | Inferior frontal gyrus pars triangularis | L | 4577 | -50 | 24 | -2 | 4.83 | .001 |
|  | Supramarginal gyrus | L | 3094 | -58 | -46 | 26 | 4.33 | .006 |
|  | Medial superior frontal gyrus | L | 3155 | -4 | 28 | 44 | 4.03 | .005 |
| Interaction: Control>Patient |  |  |  |  |  |  |  |  |
| SS>NS | N. S. |  |  |  |  |  |  |  |
| SG>NG | Anterior cingulate cortex | R | 2890 | 16 | 30 | 24 | 3.04 | .007 |
|  | Medial superior frontal gyrus | R |  | 16 | 48 | 6 | 2.95 |  |
| SB>NB | N. S. |  |  |  |  |  |  |  |

For accuracy, we observed three main effects despite no interaction with GROUP: Patients were significantly less accurate than healthy controls (F_(1,33)_ = 5.18, p = .03). Non-social videos were judged more accurately than social videos (F_(1,33)_ = 19.75, p = .0001). Additionally, accuracy between the three modalities was different (F_(2,66)_ = 14.53, p=.00001). Post-hoc pairwise t-test showed that the accuracy for the bimodal conditions were highest (vs. gesture, t = 5.81, p = .000001; vs. speech, t = 2.77, p = .007). The accuracy for the speech conditions was also higher than the gesture conditions (t = 2.14, p = .035).

For reaction time, there were also three main effects and no interaction with GROUP: patients were generally slower than healthy controls (F_(1,33)_ = 19.64, p=.00001). Reaction times for non-social content were faster than social content (F_(1,33)_ = 8.63, p=.005). Additionally, reaction times for the three modalities were different (F_(2,66)_ = 10.14, p=.0001). Post-hoc pairwise t-test showed that the reaction time for gesture modality was significantly slower than the other modalities (|t|_min_ = 4.03, p_max_ = .0001), and there was no significant difference between the bimodal and the speech modalities (t = 0.21, p = .82).

The results from the behavioral task showed that patients with schizophrenia were generally slower and less accurate in the content judgement task. Additionally, the responses for non-social videos were faster and more accurate. With regard to modality, responses in the bimodal conditions were the most accurate and the fastest.

### 3.2 fMRI results

#### 3.2.1 Social-abstract (S) > Non-social-concrete (N)

We report whole-brain fMRI results for S > N comparisons in **Figure 2** and **Table 2**. For the speech conditions (SS > NS), healthy controls activated an extensive fronto-temporal-parietal network including the bilateral inferior frontal gyrus (IFG) and the temporal lobe, the dorsolateral prefrontal cortex (dlPFC) and mPFC, and the left supramarginal gyrus; patients revealed similar regions for this comparison, and we observed no group difference for social > non-social speech. For the gesture conditions (SG > NG), controls activated the bilateral PFC and IFG; patients activated the bilateral prefrontal cortex. Group interaction (Control (SG > NG) > Patient (SG > NG)) suggests that patients showed reduced activation in the mPFC and the anterior cingulate cortex for the social gesture condition when compared to controls (**Figure 2B**). In the bimodal condition (SB > NB), both controls and patients activated regions similar to that of the speech condition. For patients, we additionally reported modality*content interaction in the **Supplement S2**, which shows that patients’ aberrant processing of social gestures is improved in the bimodal modality.

**Figure 2.**
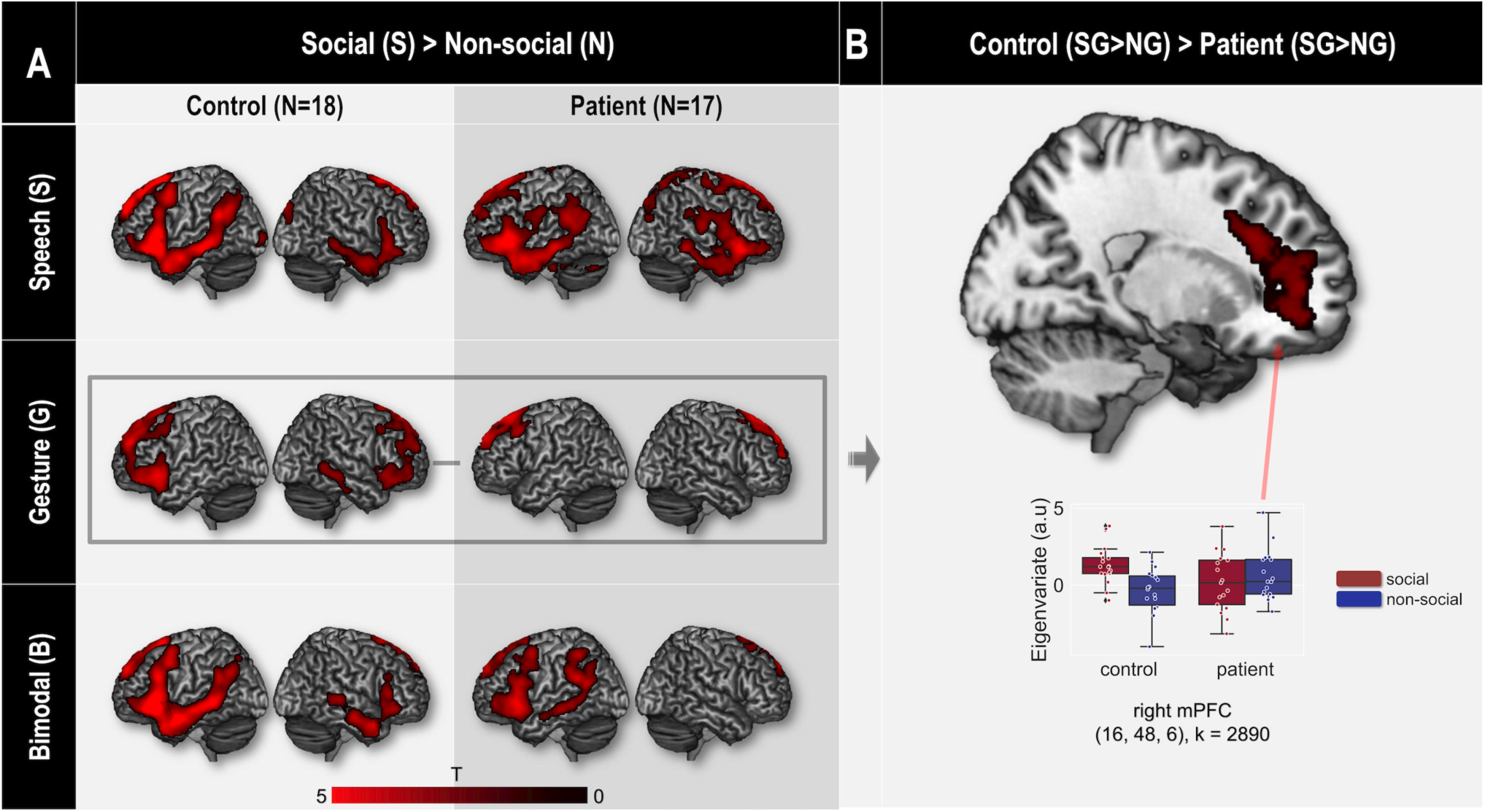
Activation maps for social-abstract > non-social-concrete videos (S > N). Panel A: S > N contrasts within each modality (S: Speech, G: Gesture, B: bimodal) for controls and patients. Panel B: interaction analysis (Control > Patient) in the gesture modality (SG > NG) together with box- and swarm-plots of eigenvariates for selected clusters. All results are correct for multiple comparison with Monte-Carlo simulation with k > 2268 voxels.

#### 3.2.2. Non-social-concrete (N) > Social-abstract (S)

We report whole-brain fMRI results for N > S comparisons in **Figure 3** and **Table 3**. For the speech conditions (NS > SS), healthy controls activated the left pre/postcentral gyrus, supramarginal gyrus, and the left insula, whereas patients did not reveal any significant activations for this comparison. The group interaction (Control (NS > SS) > Patient (NS > SS)) suggests that, when compared to controls, patients showed reduced activation in the left postcentral gyrus and the right insula for the processing of non-social content in the speech-only modality (**see Figure 3B**). For gesture conditions (NG > SG), controls showed increased activation for the non-social content in the bilateral posterior temporal gyrus, supramarginal gyrus, and occipital cortices, as well as the left pre/postcentral gyrus and the left insula. Patients also activated the bilateral posterior temporal gyrus and occipital lobe, as well as the left pre/postcentral gyrus. The group interaction revealed no significant clusters. For bimodal conditions (NB > SB), both controls and patients activated regions that are comparable to that of the gesture conditions. Additionally, for patients, bimodal input seems to enhance their aberrant processing of non-social speech, as reported in **Supplement S2**.

**Table 3:** Peak MNI coordinates of within- and between-group significant brain comparisons for non-social > social conditions. Cluster-level uncorrected p valeue is provided for each cluster, asterisk indicates signifance (p < 0.05) with cluster-level FWE correction.

| Contrasts | Anatomical Region (Peak) | Hem. | k | Peak MNI coordinates |  |  | t | p (unc.) |
| --- | --- | --- | --- | --- | --- | --- | --- | --- |
|  |  |  |  | x | y | z |  |  |
| Control |  |  |  |  |  |  |  |  |
| NS>SS | Middle cingulate cortex | L | 17667 | -48 | -28 | 42 | 4.18 | <.001* |
| NG>SG | Middle Occipital gyrus | L | 18822 | -46 | -72 | 4 | 6.48 | <.001* |
|  | Middle temporal gyrus | R | 2620 | 50 | -64 | 8 | 5.01 | .01 |
| NB>SB | Middle temporal gyrus | L | 15424 | -50 | -72 | 6 | 4.55 | <.001* |
|  | Supplementary motor area | R | 8910 | 6 | -12 | 56 | 3.7 | <.001* |
| Patient |  |  |  |  |  |  |  |  |
| NS>SS | N. S. |  |  |  |  |  |  |  |
| NG>SG | Middle temporal gyrus | L | 49718 | -48 | -70 | 6 | 6.9 | <.001* |
| NB>SB | Middle occipital gyrus | L | 22719 | -30 | -88 | 2 | 4.86 | <.001* |
|  | Olfactory cortex | L | 3347 | -8 | 26 | -4 | 4.2 | .004 |
| Interaction: Control>Patient |  |  |  |  |  |  |  |  |
| NS>SS | Insula | R | 2550 | 36 | -12 | 2 | 3.05 | .011 |
|  | Postcentral gyrus | L | 3630 | -66 | -14 | 22 | 2.92 | .003 |
| NG>SG | N. S. |  |  |  |  |  |  |  |
| NB>SB | N. S. |  |  |  |  |  |  |  |

**Figure 3.**
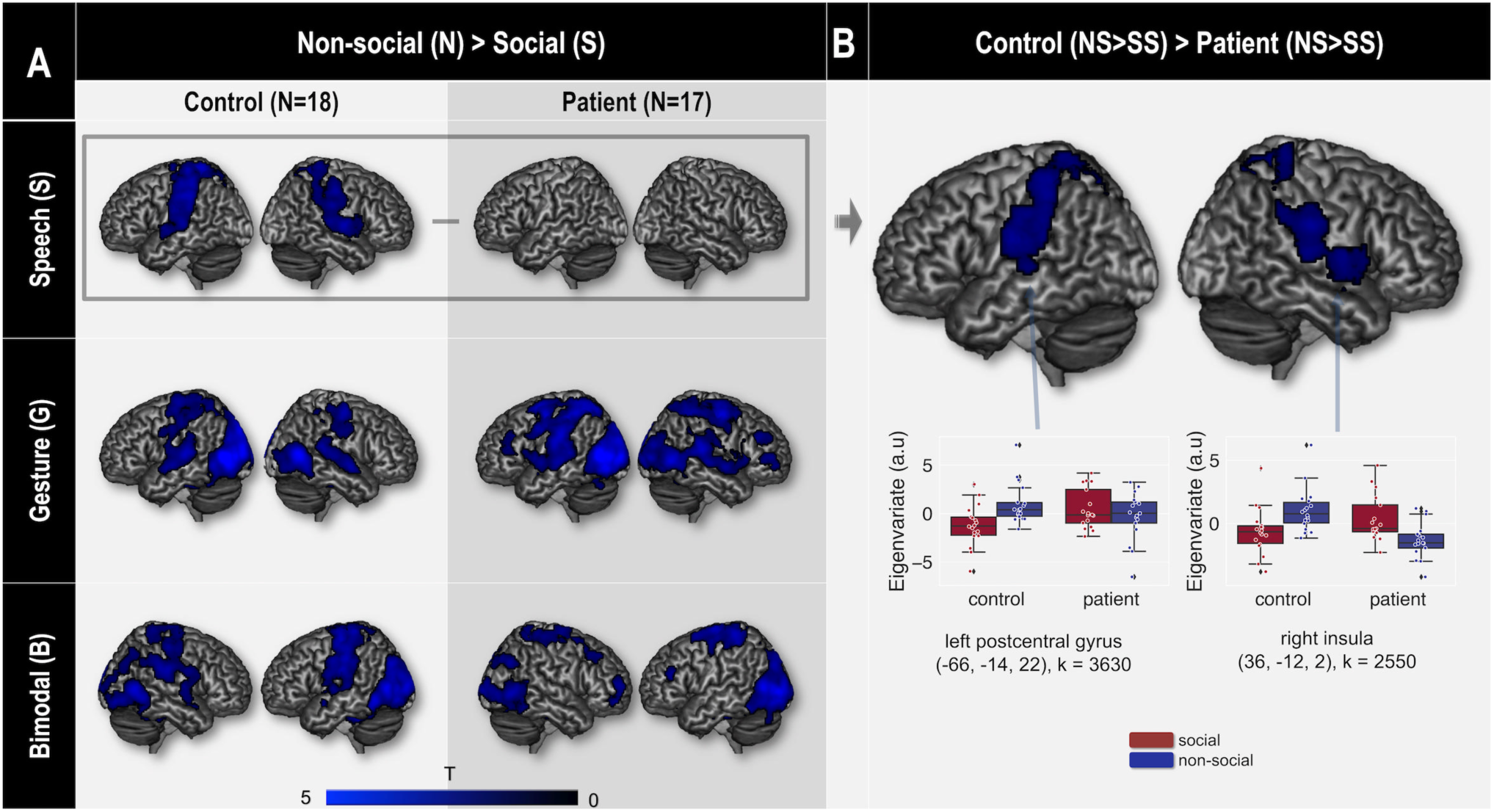
Activation maps for non-social-concrete > social-abstract videos (N > S). Panel A: N > S contrasts within each modality (S: Speech, G: Gesture, B: Bimodal) for controls and patients. Panel B: interaction analysis (Control > Patient) in the speech modality (NS > SS) together with box- and swarm-plots of eigenvariates for selected clusters. All results are correct for multiple comparison with Monte-Carlo simulation with k > 2268 voxels.

### 3.3 Exploratory correlation analyses

In patients, for the NS condition, we found that the accuracy for the NS condition correlate negatively with the SANS composite scores of the patients (r = -0.52, p = 0.03, power = 0.63; **Figure 4A**). Additionally, SANS 1 (flat affect) and SANS 2 (alogia) scores correlate negatively with the accuracy for the NS condition (SANS 1: r = -0.62, p = 0.008, power = 0.79, **Figure 4B**; SANS 2: r = -0.63, p = 0.006, power = 0.82, **Figure 4C)**.

**Figure 4.**
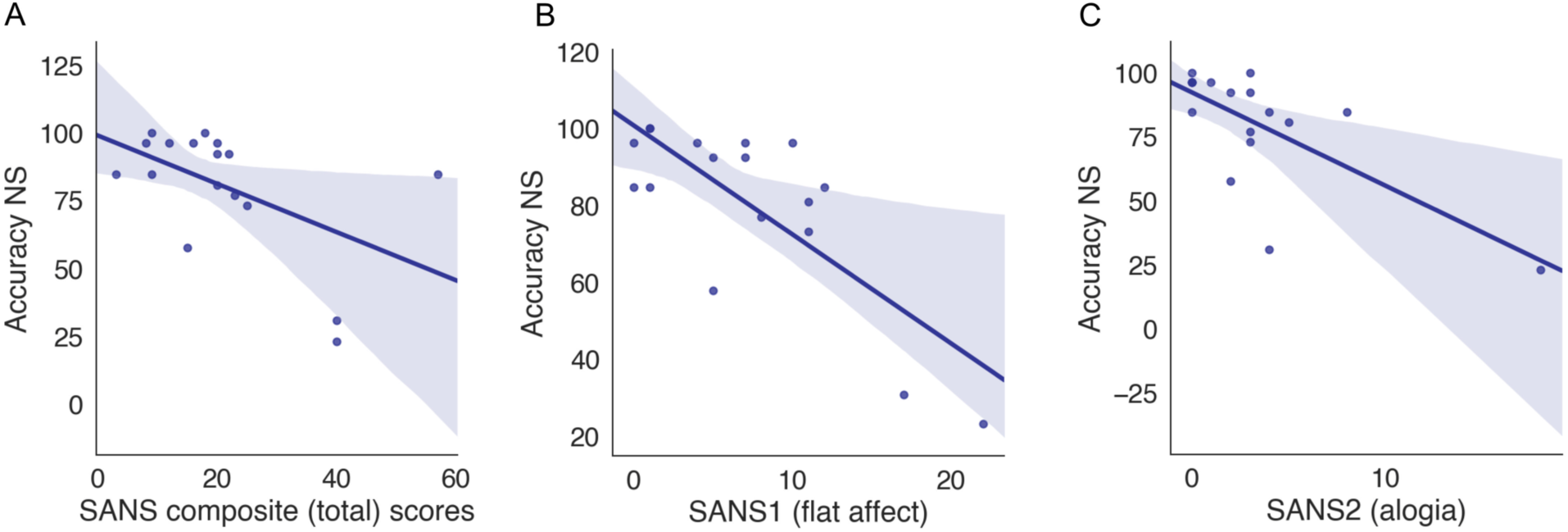
Significant negative correlations between patients’ accuracy for the NS condition and A) patients’ SANS composite scores, B) their SANS 1 (flat affect) scores, and C) their SANS 2 (alogia) scores.

## 4. Discussion

With an fMRI study on the perception of social-abstract and non-social-concrete stimuli in auditory-speech, visuo-gestural, and bimodal modalities, we observed modality-specific neural modulation in schizophrenia: In comparison to controls, reduced neural activity in patients was observed only for social gesture and non-social speech. Moreover, in the bimodal condition, neural activation for both social and non-social contents was comparable in patients and controls.

### 4.1 Social information processing in schizophrenia

In the current study, patients showed dissociable neural modulation during the processing of social content in speech and gesture modalities. In the speech modality, both controls and patients activated a left-lateralized set of brain regions, including the dlPFC, mPFC, the IFG, the temporal lobe, and the angular/supramarginal gyrus, without any group difference. This finding replicates results from our previous study showing supramodal social-abstract processing of healthy individuals (Straube et al., 2013b), and is consistent with earlier studies in basic research on the role of the mPFC in both perceiving social-related stimuli and mentalizing social intentions (Mitchell et al., 2005; Mitchell et al., 2002; Van Overwalle, 2009). The observed left IFG and temporal lobe activation is also in line with the literature on the neural substrates of abstract vs. concrete semantics (Binder et al., 2009; Binder et al., 2005), as the videos in the social-abstract condition, irrespective of modality, are more abstract than the non-social, object-related condition. The fact that we did not find any group differences in social-abstract speech processing suggests that patients with schizophrenia exhibit intact neural processing of social content presented in this modality. This finding complies with a previous language study in schizophrenia, in which patients also activated a comparable left fronto-temporal network to controls when they processed abstract vs. concrete visual sentences (Kuperberg et al., 2008). Together, although schizophrenia is well-known for its social cognition deficits (Green et al., 2015), as well as impairments in the perception of affective face or pitch (Edwards et al., 2001; Leitman et al., 2005; Morris et al., 2009), patient’s processing of social speech may remain intact. In the gesture modality, however, although patients activated the mPFC for the social vs. non-social stimuli, this activation was reduced when compared to controls. Notably, such modality-specific neural modulation is, for the first time, reported for social information processing in schizophrenia. Notably, in addition to the mPFC modulation, in schizophrenia, we did not observe the activation for social gestures in the left IFG. The left IFG has been linked to motor planning of gesture (Walther et al., 2013); and, together with the DLPFC, left IFG activation has been shown to be modulated in schizophrenia for social gesture planning (Stegmayer et al., 2018). Thus, our results, from a perception perspective, corroborates these extant studies, showing that both the PFC and the left IFG functional modulation may give rise to gesture-specific social perception deficits.

### 4.2 Non-social information processing in schizophrenia

For the processing of non-social (object-related) information, again, the neural modulation in patients showed an apparent dissociation. In the gesture modality, both controls and patients activated the bilateral occipital-parietal cortices, STG, LOTC, insula, and the pre/postcentral gyrus. In the speech modality, although these regions were similarly activated in controls, their brain activation was significantly reduced for patients. The group comparison suggests reduced activation in schizophrenia patients in the left insula and the left postcentral gyrus for non-social speech. Of note, the observed regions for non-social and object-related information processing overlap with regions considered as part of the putative mirror neuron network, which is not only important for action observation and imitation, but also for the understanding of object- and motor-related features in verbal form (Buccino et al., 2001; Di Pellegrino et al., 1992; Iacoboni et al., 1999; Rizzolatti & Craighero, 2004). This process would require mental simulation of sensorimotor experience (Barsalou, 2008; Pulvermüller, 2005; Pulvermüller et al., 2005). Additionally, the LOTC is also crucially involved in the perception of biological motion, object, as well as tool-use (Bi et al., 2016; Higuchi et al., 2007; Lingnau & Downing, 2015). Moreover, these regions are also reported to support the processing of concrete linguistic information (Binder et al., 2009; Binder et al., 2005). Our data from the control group suggest that these regions support the processing of non-social-concrete features, irrespective of encoding modality. This finding is in line with the embodiment view of action and language processing (Barsalou, 2008; De Stefani & De Marco, 2019). With regard to the patients, we observed normal neural processing of non-social content in the gesture modality, supporting a previous study (Horan et al., 2014), which reported intact mirror neuron activity in schizophrenia (but see (Thakkar et al., 2014)). However, as we also observed reduced bilateral postcentral gyrus and right insula activity for patients for non-social speech, in turn, this would imply that motor simulation, as required for processing object-related features from auditory speech, might still be impaired in schizophrenia (Kuperberg et al., 2008; Thakkar et al., 2014). This impairment concurs with the reported deficits of schizophrenia in action imitation (Matthews et al., 2013; Mehta et al., 2014; Park et al., 2008; Thakkar et al., 2014; Walther et al., 2015; Walther et al., 2013), where certain degrees of motor simulation is required. However, adding to prior research on schizophrenia’s motor impairments, our results provide a more nuanced version: Motor-related semantics as delivered by pantomime does not necessarily lead to reduced neural activation in schizophrenia; rather, modulated neural activity is only observed for speech contents simulating these concrete motor-actions. This might be suggesting that patients with schizophrenia are impaired in a mirror-mechanism simulating concrete-motor events with speech.

In the NS condition, we also observed negative correlation between patients’ SANS composite and subscores and their task accuracy. This evidence converges with previous research, corroborating the potential role of the mirror neuron system during embodiment of non-social information (e.g., action imitation and observation), as well as its relation to the development and persistence of negative symptoms (Buccino & Amore, 2008; Mehta et al., 2014). Our correlational finding, although being exploratory (see 4.4), links the mirror neuron system to the reported studies showing correlation between compromised gesture performance and more severe negative symptoms (Park et al., 2008; Walther et al., 2019), tentatively suggests that the theoretical link between gesture and negative symptom may partially derive from motor-simulation. Notably, in the current study, for social processing, we did no observe any correlations between brain activations / behavior and symptom measures, especially positive symptoms such as hallucinations, and positive formal thought disorders, as in Straube et al. (2014). Clearly, the exact relationship between social/non-social information processing and major symptoms of schizophrenia needs to be addressed by further research with larger samples.

### 4.3 Enhancing modality-specific information processing deficits with bimodal input

The novelty of our findings lies in the dissociable modality-specificity concerning dysfunctional neural processing of social and non-social features. Social-abstract and non-social-concrete features are functionally and neurally dissociable at the representational level (Binder et al., 2009; Mitchell et al., 2005; Mitchell et al., 2002). Besides, they might be differentially processed through either linguistic (speech) or non-linguistic (gesture) channels. It has been proposed that social-abstract concepts may be *preferentially* represented in speech, and that non-social concrete concepts are *preferentially* delivered in hand action and gesture (Paivio, 2010; Perlovsky & Ilin, 2013). Despite this theoretical proposal, however, during comprehension, healthy participants are able to process both types of information in a supramodal manner (e.g., semantic processing with unitary core systems, irrespective of encoding modality, as in (Pulvermüller et al., 2005; Straube et al., 2013b)). For patients with schizophrenia, as they exhibit similar neural activations when processing social speech and non-social gestures to controls, this might be an indication that they are at least intact in processing these contents through a *preferred* modality at representational level. But, they might show activation reduction in relevant regions when these features are conveyed in a *non-preferred* modality, as the processing of these features would require some form of mental simulation: In the case non-social information, patients are impaired in the simulation of motor-related experience from action to language (Pulvermüller et al., 2005); In contrast, when patients are presented with social information, they might be impaired when simulating social features encoded by hand gestures (but not with speech), as shown in their reduced mPFC activation. This observed modality-specific processing deficit might also suggest that patients, unlike controls, are not capable of processing social/non-social information in a supramodal manner like healthy participants, as reported in previous studies (Straube et al., 2012; Straube et al., 2013b). More importantly, extending previous studies on aberrant processing of social/non-social content in schizophrenia, our results indicate that this neural deficit is not universally present for either a specific modality or content, but rather appears only in specific combinations of these two factors.

Despite reduced neural processing of both social and non-social content in gesture and speech modalities, patients displayed intact neural processing of these features, as well as improved task accuracy in the bimodal conditions. This enhancement effect concurs with a line of proposals (Holler & Levinson, 2019), who argue for a bi-directional facilitative relation between speech and gesture (for empirical evidence, see (Cuevas et al., 2019; Drijvers et al., 2018; He et al., 2018b; Krahmer & Swerts, 2007; Wang & Chu, 2013)). More importantly, our finding extends previous basic research, suggesting the translational implication of this mechanism. In schizophrenia research, the past decade has witnessed substantial progress in the development of social cognitive training in schizophrenia (Kurtz et al., 2016; Kurtz & Richardson, 2011), with recent innovation regarding the incorporation of social stimuli from a broader range of modalities (Nahum et al., 2014). Our findings extend these approaches, proposing potential therapeutic implications of deploying naturalistic and multimodal stimuli during social cognitive training (Riedl et al., 2020), as they might be able to normalize processing of both social and non-social information, at least at a neural level. Future research is expected to further explore whether the neural enhancements can be linked to functional outcome after social cognitive training in a multimodal setting.

### 4.4 Limitations

Despite new insights, our study is limited in several aspects. Firstly, the sample size of this study (n = 17 and 18 for each group) falls within the required minimum of this type of studies (Friston, 2013). Hence, due to sample size, the patient group was limited to medicated chronic patients, and we were thus unable to test if the neural pattern is comparable to first-episode schizophrenia (Kindler et al., 2019), and were unable to attribute the findings to schizophrenia without separating out the effects of medication. Secondly, due to the limited sample size and the exploratory nature, the findings resulted from the correlation analyses needs to be treated with caution. Lastly, we did not observe any reliable relationship between BOLD signal and behavioral results: More careful, and neurobiologically plausible behavioral tasks on the social/non-social information processing are necessary for future experiments, so that their processing deficits in schizophrenia can be better examined by clinical neuroscientists and practitioners.

### 4.5 Conclusion

Here, in an fMRI study, we for the first time showed modality-specific neural modulation in schizophrenia when patients process social-abstract and non-social-concrete content in speech and gesture. Moreover, these deficits could be compensated when both speech and gesture were presented together. Our findings provide novel insights on dysfunctional multimodal communication in schizophrenia, and suggest potential therapeutic implications of employing multimodal social cognitive intervention in schizophrenia.

## 5. Acknowledgements

This research project is supported by a grant from the ‘Von-Behring-Röntgen-Stiftung’ (project no. 59-0002 and 64-0001) and by the ‘Deutsche Forschungsgemeinschaft’ (project no. DFG: STR1146/11-2 & KI588/6-2 and CRC/TRR 135/2 project A3). The study was also supported by the Core Facility Brain Imaging, Faculty of Medicine, University of Marburg, Rudolf-Bultmann-Str. 9, 35039, Marburg, Germany. The manuscript has been published as a preprint on bioRxiv.

