## Supplement for "Modality-specific dysfunctional neural processing of social-abstract and non-social-concrete information in schizophrenia"

*S1: TABLE S1*

**Table S1**. Symptom subscores, and neuropsychological measures for patients and healthy controls

| | Patients (n=17) | Controls (n=18) |
|---|---|---|
| WFT (animal) sum | 21.23 (5.33) | 21.72 (6.02) |
| WFT (P) sum | 4.26 (0.17) | 3.63 (0.33) |
| *WFT (changing category) sum | 3.28 (0.17) | 2.76 (0.39) |
| BAG subscores | 3.27 (0.58) | 3.37 (0.46) |
| BAG I (perception) | 3.65 (0.83) | 3.86 (0.58) |
| BAG II (production) | 2.97 (0.87) | 2.99 (0.79) |
| BAG III (social production/help) | 3.56 (0.93) | 3.94 (0.59) |
| BAG IV (social perception) | 2.82 (1.06) | 2.55 (0.97) |
| SAPS (composite) total | 22.35 (18.08) | |
| SAPS 1 (hallucinations) | 4.64 (6.33) | |
| SAPS 2 (Delusions) | 10.47 (9.37) | |
| SAPS 3 (Bizarre behavior) | 1.88 (2.93) | |
| SAPS 4 (Thought disorder) | 5.41 (7.00) | |
| SANS (composite) total | 21.00 (13.63) | |
| SANS 1 (Flat affect) | 7.18 (6.20) | |
| SANS 2 (Alogia) | 3.29 (4.37) | |
| SANS 3 (Apathy) | 4.76 (4.51) | |
| SANS 4 (Anhedonia) | 4.35 (4.44) | |
| SANS 5 (Attention) | 1.59 (2.58) | |

Values are presented as mean (SD). WFT: word fluency test; BAG: Brief Assessment of Gesture. Asterisk * indicates significant difference between controls and patients ($p<0.05$, two-tail t-test).

*S2. Bimodal enhancement effects within patients*

For the bimodal conditions, group interaction revealed no significant effects for the comparison between social and non-social conditions in both directions. This might suggest that bimodal social and non-social videos are normally perceived for patients with schizophrenia. Therefore, we investigated whether bimodal input normalizes aberrant social/non-social information processing for patients. Firstly, we tested for patients with schizophrenia, whether the potential normalization effects result from enhanced social/non-social processing for bimodal vs. impaired modalities (e.g., social gestures and non-social speech). We observed that, for patients, for the processing of social videos, bimodal > gesture interaction [(SB > NB) > (SG > BG)] additionally activated the bilateral IFG (**Figure S1A, Table S2**). For the processing of non-social content, bimodal > speech comparison [(NB > SB) > (NS > SS)] revealed increased brain activation in the left thalamus, the right cingulate cortex, the left occipital lobe, and the left postcentral/precentral gyrus (**Figure S1B**, **Table S2**). However, when we performed conjunction analyses between these contrasts and the contrast image from the three-way interaction analysis, no significant brain regions survived our threshold. Thus, these results do not speak for a potential normalization effect of the bimodal condition; however, the results indicate that, at least for patients, bimodal stimuli enhance social and non-social processing when compared to aberrantly processed modality.

*Figure S1*

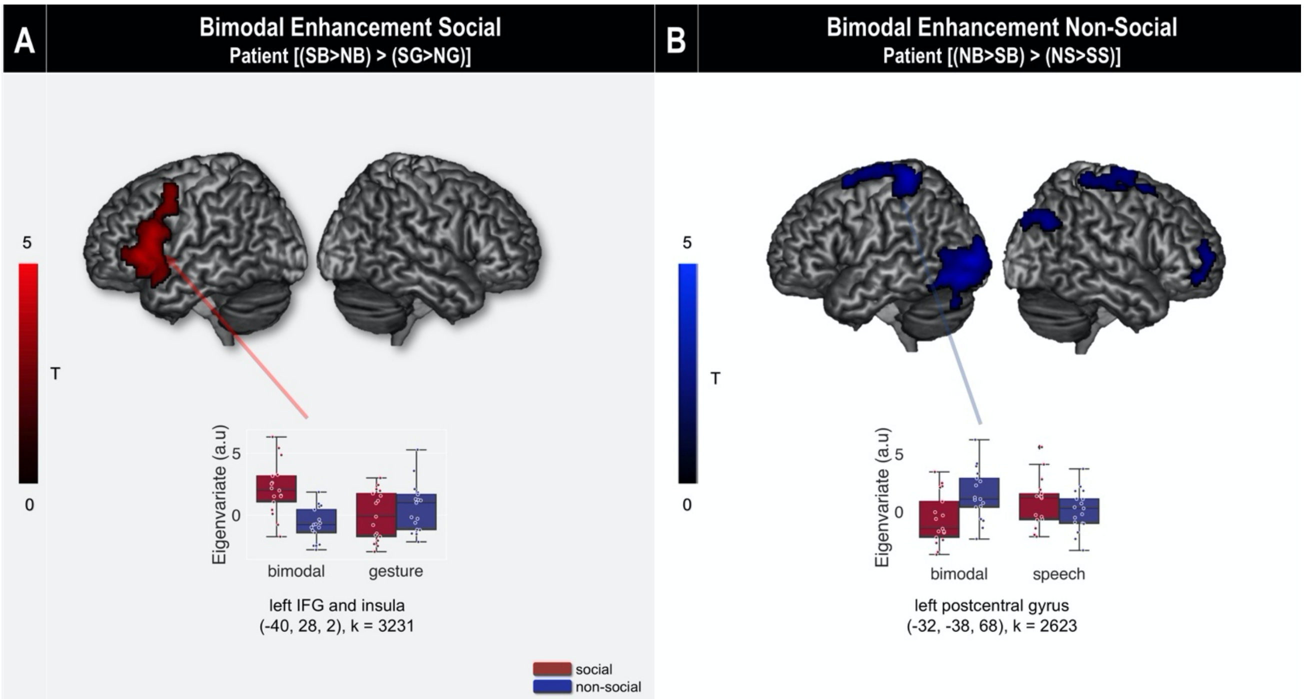

**Figure S1.** Activation maps showing bimodal normalization effect for both social (Panel A) and non-social (Panel B) conditions in patients with SZ. Panel A: Bimodal > gesture interaction for the social condition [(SB > NB) > (SG > BG)]; Panel B: Bimodal > speech interaction for the non-social condition [(NB > SB) > (NS > SS)]. Both panels are plotted together with box- and swarm-plots of eigenvariates for selected clusters. All results are correct for multiple comparison with Monte-Carlo simulation with k > 2268 voxels.

Table S2

**Table S2**: Peak MNI coordinates of modality*content interactions within the patient group (bimodal enhancement effect). Cluster-level uncorrected p valeue is provided for each cluster, asterisk indicates significance ($p < 0.05$) with cluster-level FWE correction.

| | | | | Peak MNI coordinates | | | | |
|---|---|---|---|---|---|---|---|---|
| Contrasts | Anatomical Region (Peak) | Hem. | k | x | y | z | t | p (unc.) |
| **(SB>NB)>(SG>BG)** | Inferior frontal gyrus pars triangularis | L | 3231 | -40 | 28 | 2 | 3.84 | .005 |
| **(NB>SB)>(NS>SS)** | Olfactory gyrus | L | 2446 | -8 | 26 | -4 | 3.7 | .012 |
| | Middle occipital gyrus | L | 2786 | -30 | -90 | 2 | 3.7 | .008 |
| | Middle cingulate cortex | R | 2782 | 10 | -42 | 34 | 3.65 | .008 |
| | Postcentral gyrus | L | 2623 | -32 | -38 | 68 | 3.48 | .010 |
